# The mRNA decapping machinery targets *LBD3/ASL9* to mediate apical hook and lateral root development in *Arabidopsis*

**DOI:** 10.1101/2022.07.06.499076

**Authors:** Zhangli Zuo, Milena Edna Roux, Jonathan Renaud Chevalier, Yasin F. Dagdas, Takafumi Yamashino, Søren Diers Højgaard, Emilie Knight, Lars Østergaard, Eleazar Rodriguez, Morten Petersen

## Abstract

Multicellular organisms perceive and transduce multiple cues to optimize development. Key transcription factors drive developmental changes, but RNA processing also contributes to tissue development. Here, we report that multiple decapping deficient mutants share developmental defects in apical hook, primary and lateral root growth. More specifically, *LATERAL ORGAN BOUNDARIES DOMAIN 3* (*LBD3*)/*ASYMMETRIC LEAVES 2-LIKE 9* (*ASL9*) transcripts accumulate in decapping deficient plants and can be found in complexes with decapping components. Accumulation of *ASL9* inhibits apical hook, primary root growth and lateral root formation. Interestingly, exogenous auxin application restores lateral roots formation in both *ASL9* over-expressors and mRNA decay-deficient mutants. Likewise, mutations in the cytokinin transcription factors type-B ARABIDOPSIS RESPONSE REGULATORS (B-ARRs) *ARR10* and *ARR12* restore the developmental defects caused by over-accumulation of capped *ASL9* transcript upon *ASL9* overexpression. Most importantly, loss-of-function of *asl9* partially restores apical hook and lateral root formation in decapping deficient mutants. Thus, the mRNA decay machinery directly targets *ASL9* transcripts for decay, possibly to interfere with cytokinin/auxin responses, during development.

## Introduction

Understanding proper tissue development requires information about diverse cellular mechanisms controlling gene expression. Much work has focused on of the transcriptional networks that govern stem cell differentiation. For example, ectopic expression of *LATERAL ORGAN BOUNDARIES DOMAIN (LBD)/ASYMMETRIC LEAVES2-LIKE (ASL*) genes is sufficient to induce spontaneous proliferation of pluripotent cell masses in plants, a reprogramming process triggered *in vitro* by complementary/Yin-Yang phytohormones auxin and cytokinin (Fan et al., 2012; Schaller et al., 2015). Auxin and cytokinin responses are essential for a vast number of developmental processes in plants including post-embryonic reprograming and formation of the apical hook to protect the meristem during germination in darkness (Chaudhury et al., 1993; Hu et al., 2017) as well as lateral root (LR) formation (Jing and Strader, 2019). Loss-of-function mutants in genes that regulate auxin-dependent transcription such as *auxin-resistant1* (*axr1*) exhibit defective hooking and LR formation (Estelle and Somerville, 1987; Lehman et al., 1996). In addition, type-B ARABIDOPSIS RESPONSE REGULATORS (B-ARRs) ARR1, ARR10 and ARR12 work redundantly as transcriptional activators to regulate cytokinin targets including type-A ARRs, which are negative regulators of cytokinin signaling in shoot development and LR formation (Riefler et al., 2006; Ishida et al., 2008; Xie et al., 2018). Exogenous cytokinin application disrupts LR initiation by blocking pericycle founder cell transition from G2 to M phase (Li et al., 2006). Thus, reshaping the levels of certain genes leads to changes in cellular identity. As developmental programming must be tightly regulated to prevent spurious development, the expression of these transcription factors may be controlled at multiple levels (Tatapudy et al., 2017). However, most developmental studies focus on their transcription rates and overlook the contribution of mRNA stability or decay to these events (Crisp et al., 2016).

Eukaryotic mRNAs contain stability determinants including the 5’ 7-methylguanosine triphosphate cap (m7G) and the 3’ poly-(A) tail. mRNA decay is initiated by deadenylation, followed by degradation via either 3’-5’ exosomal exonucleases and SUPPRESSOR OF VCS (SOV)/DIS3L2 or via the 5’-3’ exoribonuclease activity of the decapping complex (Garneau et al., 2007; Sorenson et al., 2018). This complex includes the decapping holoenzyme composed of the catalytic subunit Decapping 2 (DCP2) and its cofactor DCP1 along with other factors (DCP5, DHH1, VCS, LSM1-7 complex and PAT1), and the Exoribonuclease (XRN) that degrades monophosphorylated mRNA. As a central platform, PAT1 (Protein Associated with Topoisomerase II, PAT1b in mammals) forms a heterooctameric complex with LSM(Like-sm)1-7 at 3’ end of a mRNA to engage transcripts containing deadenylated tails thereafter recruits other decapping factors and interacts with them using different regions, these decapping complex and mRNAs can aggregate into distinct cytoplasmic foci called processing bodies (PBs) (Brengues et al., 2005; Balagopal and Parker, 2009;Ozgur et al., 2010; Chowdhury et al., 2014; Charenton et al., 2017; Lobel et al., 2019). Beyond *DCP* genes, deletion of *PAT1* gene in yeast exhibits the strongest temperature sensitive phenotype compared to other decapping factors genes (Bonnerot et al., 2000).

mRNA decay regulates mRNA levels and thereby impacts cellular reprogramming (Newman et al., 2017; Essig et al., 2018). We and others have shown that the decapping machinery is involved in stress and immune responses (Xu and Chua, 2012; Merret et al., 2013; Roux et al., 2015; Perea-Resa et al., 2016; Crisp et al., 2017; Yu et al., 2019), and that RNA binding proteins can target selected mRNAs for decay (Gerstberger et al., 2014; Perea-Resa et al., 2016; Yu et al., 2019). Postembryonic lethality (Xu et al., 2006) and stunted growth phenotypes (Xu and Chua, 2009; Perea-Resa et al., 2012) associated with disturbance of the decay machinery indicate the importance of mRNA decapping and decay machinery during plant development. However, while much has been learned about how mRNA decapping regulates plant stress responses (Perea-Resa et al., 2016; Yu et al., 2019; Zuo et al., 2021), far less is known about how decapping contributes to plant development.

*Arabidopsis dcp1, dcp2* and *vcs* mutants display postembryonic lethality whereas *lsm1alsm1b, pat* triple mutant and *dcp5* knock-down mutants only exhibit abnormal development (Xu et al., 2006; Xu and Chua, 2009; Perea-Resa et al., 2012; Zuo et al., 2022a; Zuo et al., 2022b). All these differences suggest that mutations in mRNA decay components may cause pleiotropic phenotypes not directly linked to mRNA decapping and decay deficiencies (Riehs-Kearnan et al., 2012; Gloggnitzer et al., 2014; Roux et al., 2015). For example, it has been proposed that lethality in some mRNA decay loss-of-function mutants is not due to decay deficiencies *per se*, but to the activation of immune receptors which evolved to surveil microbial manipulation of the decay machinery (Roux et al., 2015). In line with this, loss-of-function of *AtPAT1* inappropriately triggers the immune receptor SUMM2, and *Atpat1* mutants consequently exhibit dwarfism and autoimmunity (Roux et al., 2015). Thus, PAT1 is under immune surveillance and PAT proteins are best studied in SUMM2 loss-of-function backgrounds.

Here, we studied the impact of mRNA decapping during development. For this, we have analysed 3 sequential mRNA decapping mutants *dcp2-1, dcp5-1* and *pat* triple mutant (*pat1-1path1-4path2-1summ2-8*), revealing that the mRNA decay machinery targets the important developmental regulator *ASL9*. Specifically, disruption of the mRNA decay machinery promotes *ASL9* accumulation, and this in turn contributes to inhibit apical hook and lateral root formation. Interestingly, these developmental defects, which are observed in mRNA decapping deficient mutants and *ASL9*-overexpressors, can be salvaged through disruption of cytokinin signalling or exogenous application of auxin. Importantly, mutations in *asl9* also partially restores the developmental defects including apical hook and lateral root formation in decapping mutants. These observations indicate that the mRNA decay machinery is fundamental to developmental decision making.

## Results

### mRNA decapping deficiency causes deregulation of apical hooking

We and others have reported that mutants of mRNA decay components exhibit abnormal developmental phenotypes including postembryonic death and stunted growth (Xu et al., 2006; Xu and Chua, 2009; Perea-Resa et al., 2012; Roux et al., 2015; Zuo et al.,2022b), indicating mRNA decay may be needed for proper development. To assess this, we explored readily scorable phenotypic evidence of defective development. Since apical hooking can be exaggeratedly induced by exogenous application of ethylene or its precursor ACC, we germinated seedlings in darkness in the presence or absence of ACC (Bleecker et al., 1988; Guzman and Ecker, 1990). Interestingly, all 3 sequential mRNA decapping mutants tested *dcp2-1, dcp5-1* and *pat* triple mutant were hookless and unable to make the exaggerated apical hook under ACC treatment (Fig. 1A, B, S1A&B), being that *dcp2-1* exhibit the strongest hookless phenotype. Since *dcp2-1* is postembryonic lethal, we used seeds from a parental heterozygote to score for hook formation, and subsequentially confirmed by genotyping that all hookless seedlings were *dcp2-1* homozygotes. This, and the fact that ACC treatment leads to massive increase of DCP5-GFP (Chicois et al., 2018) and Venus-PAT1(Zuo et al., 2022b) foci in hook regions (Fig.1C), all suggest that mRNA decapping is required for apical hooking.

**Fig 1.**
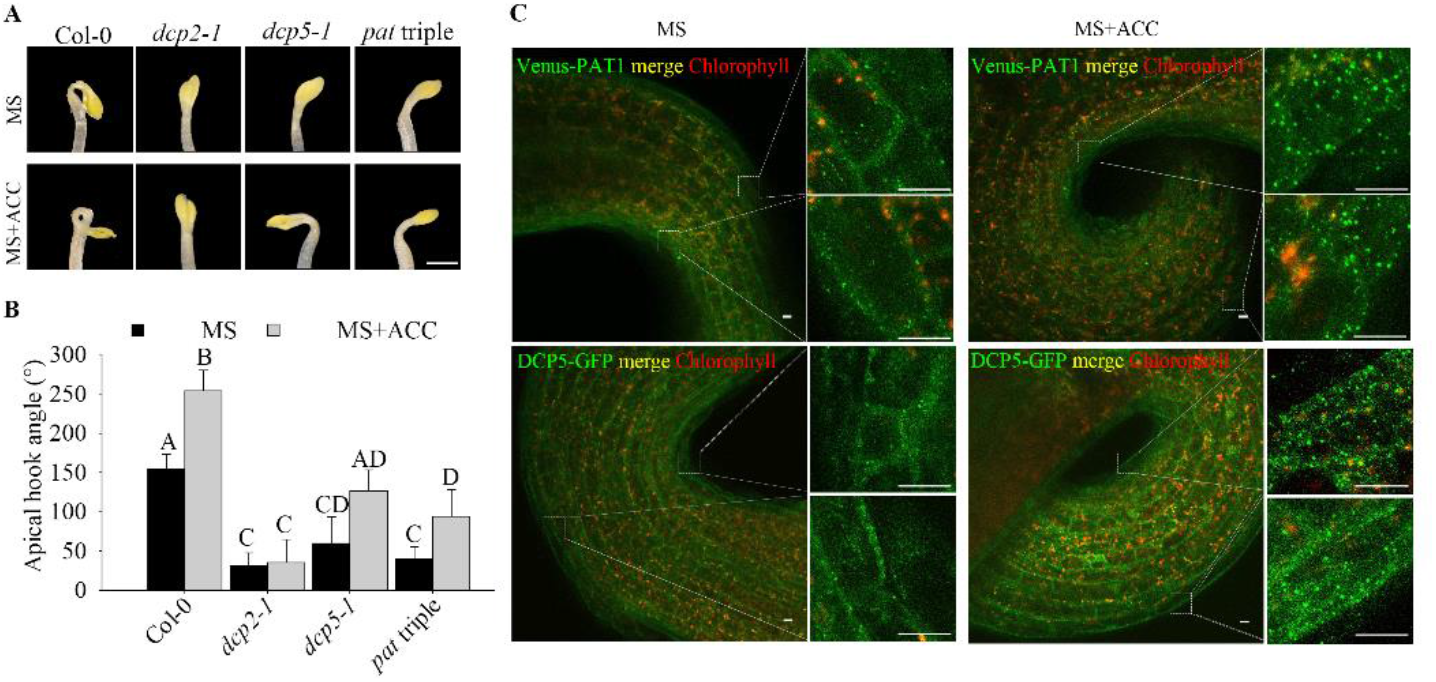
mRNA decapping deficiency causes deregulation of apical hooking. Hook phenotypes (**A**) and apical hook angles (**B**) in triple response to ACC treatment of etiolated Col-0, *dcp2-1, dcp5-1* and *pat* triple seedlings. The experiment was repeated 3 times, in each repeat sample size (n)>30 for each genotype and treatment, and representative pictures are shown. The scale bar indicates 1mm. Bars marked with the same letter are not significantly different from each other (P-value>0.05). (**C**) Representative confocal microscopy pictures of hook regions following ACC treatment. Dark-grown seedlings with either Venus-PAT1 (Top) or DCP5-GFP (Bottom) on MS or MS+ACC plates for 4 days. Scale bars indicate 10 μm.

### mRNA decay machinery targets *ASL9* for decay

To search for transcripts responsible for the hookless phenotype, we revisited our previous RNA-seq data for *pat* triple mutant (Zuo et al., 2022b) and verified that transcripts of *ASL9* (*ASYMMETRIC LEAVES 2-LIKE 9*, also named *LBD3, LOB DOMAIN-CONTAINING PROTEIN 3*) accumulated specifically in *pat* triple mutants (Zuo et al., 2022b). ASL9 belongs to the large AS2/LOB (ASYMMETRIC LEAVES 2/LATERAL ORGAN BOUNDARIES) family (Matsumura et al., 2009) which includes key regulators of organ development (Xu et al., 2016). Interestingly, the ASL9 homologue ASL4 negatively regulates brassinosteroids accumulation to limit growth in organ boundaries, and overexpression of *ASL4* impairs apical hook formation and leads to dwarfed growth (Bell et al., 2012). While *ASL4* mRNA did not accumulate in *pat* triple mutants (Data Set S1), we hypothesized that ASL9 could also interfere with apical hook formation. We therefore analyzed mRNA levels of *ASL9* in ACC-treated seedlings and verified that all 3 sequential mRNA decapping mutants accumulated up to 30-fold higher levels of *ASL9* transcript compared to ACC treated Col-0 seedlings (Fig. 2A). Concordantly, two over-expressor lines of *ASL9 Col-0/oxASL9* and *Col-0/oxASL9-VP16* (Naito et al., 2007) also exhibited hookless phenotypes (Fig. 2B&C). However, we did not observe any changes including tighter apical hooks in *asl9-1* mutants (Fig. S1C&D) suggesting other members of the AS2/LOB family act redundantly in this process. Nevertheless, these results indicate that apical hook formation in mRNA decapping deficient mutants is compromised, in part, might be due to misregulation of *ASL9*.

**Fig 2.**
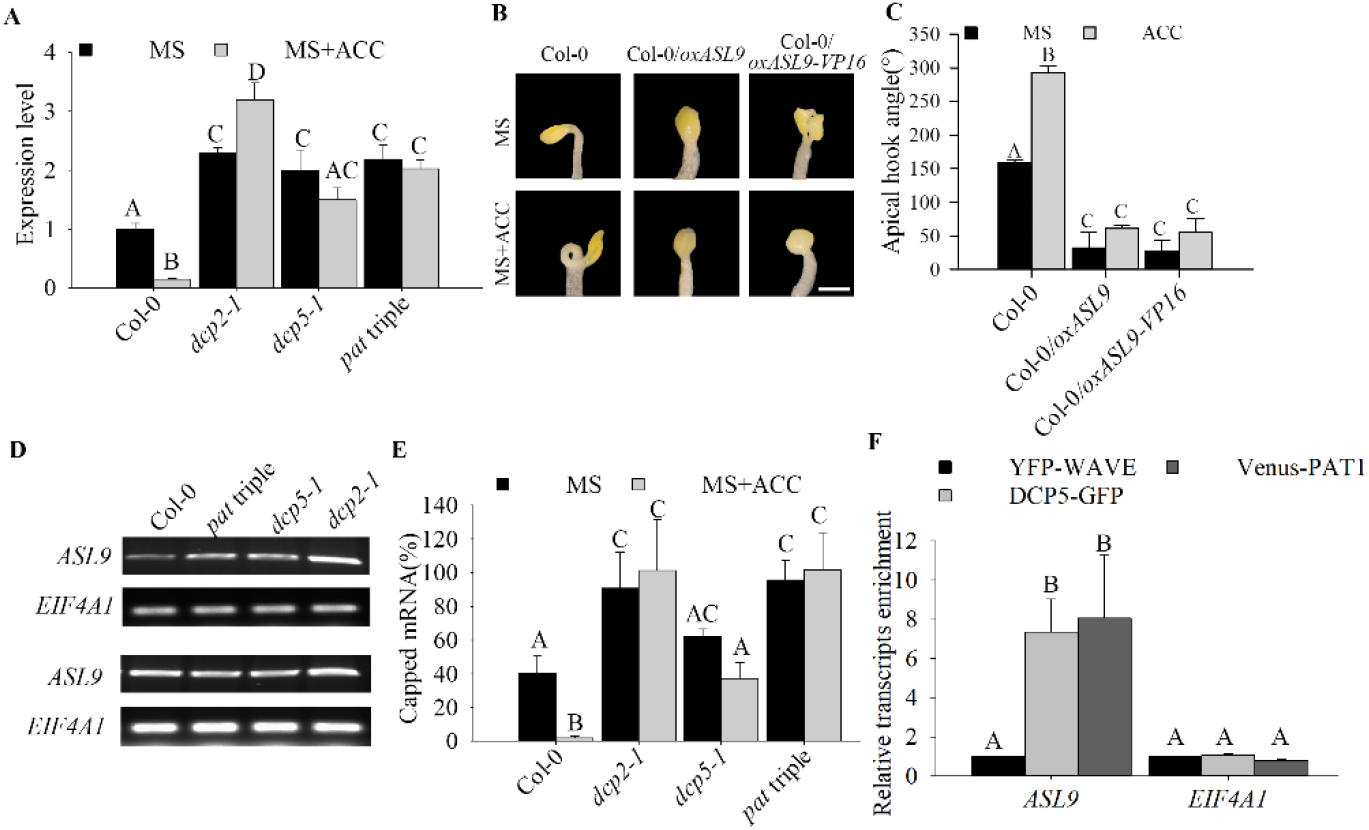
mRNA decay machinery targets *ASL9* for decay. (**A**) *ASL9* mRNA levels in cotyledons and hook regions of dark-grown Col-0, *dcp2-1, dcp5-1* and *pat* triple seedlings under control or ACC treatment. Error bars indicate SE of bio-triplicates. Hook phenotypes(**B**) and apical hook angles(**C**) of triple response to ACC treatment of etiolated seedlings of Col-0, Col-0/*oxASL9* and Col-0/*OxASL9-VP16*.The experiment was repeated 3 times, in each repeat sample sample size (n)>15 for each genotype and treatment, and representative pictures are shown. The scale bar indicates 1mm. (**D**) Accumulation of capped transcripts of *ASL9* analyzed in 4-day-old MS grown etiolated seedlings of Col-0, *pat* triple, *dcp5-1* and *dcp2-1* by 5’-RACE-PCR. RACE-PCR products obtained using a low (upper panel) and high (bottom panel) number of cycles are shown. *EIF4A1* RACE-PCR products were used as loading control. (**E**) Capped *ASL9* transcript levels using XRN1 susceptibility assay in cotyledons and hook regions of dark-grown Col-0, *dcp2-1, dcp5-1* and *pat* triple seedlings. Error bars indicate SE (n=3). (**F**) DCP5 and PAT1 bind *ASL9* transcripts. 4-day dark-grown plate seedlings with DCP5-GFP or Venus-PAT1were taken for RIP assay. *ASL9* transcript levels were normalized to those in RIP of YFP-WAVE as a non-binding control. *EIF4A1* was used as a negative control. Error bars indicate SE (n=3).

To determine whether *ASL9* is a target of the decapping complex, we performed 5’-RACE assays and found significantly higher levels of capped *ASL9* in mRNA decapping mutant seedlings than in Col-0 (Fig 2D). We also assayed for capped *ASL9* transcripts in ACC and mock-treated mRNA decapping mutants. By calculating the ratio between capped versus total *ASL9* transcripts, we verified that with ACC treatment, mRNA decapping mutants accumulated significantly higher levels of capped *ASL9* transcripts than Col-0 (Fig. 2E). Moreover, RNA immunoprecipitation (RIP) revealed enrichment of *ASL9* in DCP5-GFP and Venus-PAT1 plants compared to a MYC-YFP control line (YFP-WAVE) (Fig. 2F), indicating mRNA decapping components directly bind *ASL9* transcripts. These data confirms that *ASL9* mRNA can be found in mRNA decapping complexes, and that mRNA decapping regulates *ASL9* mRNA levels and contributes to ACC-induced apical hook formation.

### Accumulation of *ASL9* suppresses LR formation

LR formation is another example of post embryonic development. In *Arabidopsis* the first stage of LR formation requires that xylem pericyle pole cells change fate to become LR founder cells, a process positively regulated by auxin and negatively regulated by cytokinin and ethylene (Jung and McCouch, 2013; Weijers et al., 2018). We therefore examined LR formation in mRNA decapping deficient mutants *dcp5-1* and *pat* triple mutants and in both *ASL9* over-expressors and verified that LR formation was dramatically impaired in all genotypes tested (Fig. 3A, B, S2A&B). However, like seen for apical hooking, *asl9-1* also appeared to display normal LR formation (Fig. S2C&D). Nevertheless, LR formation defects in *dcp5-1* and *pat* triple mutants indicate that mRNA decapping is required for the commitment to LR formation. This is further substantiated by the fact that auxin application leads to a massive increase of DCP5-GFP and Venus-PAT1 foci in root regions (Fig 3C). Collectively, these data indicate mRNA decapping machinery, targeting *ASL9*, also contributes to LR formation.

**Fig 3.**
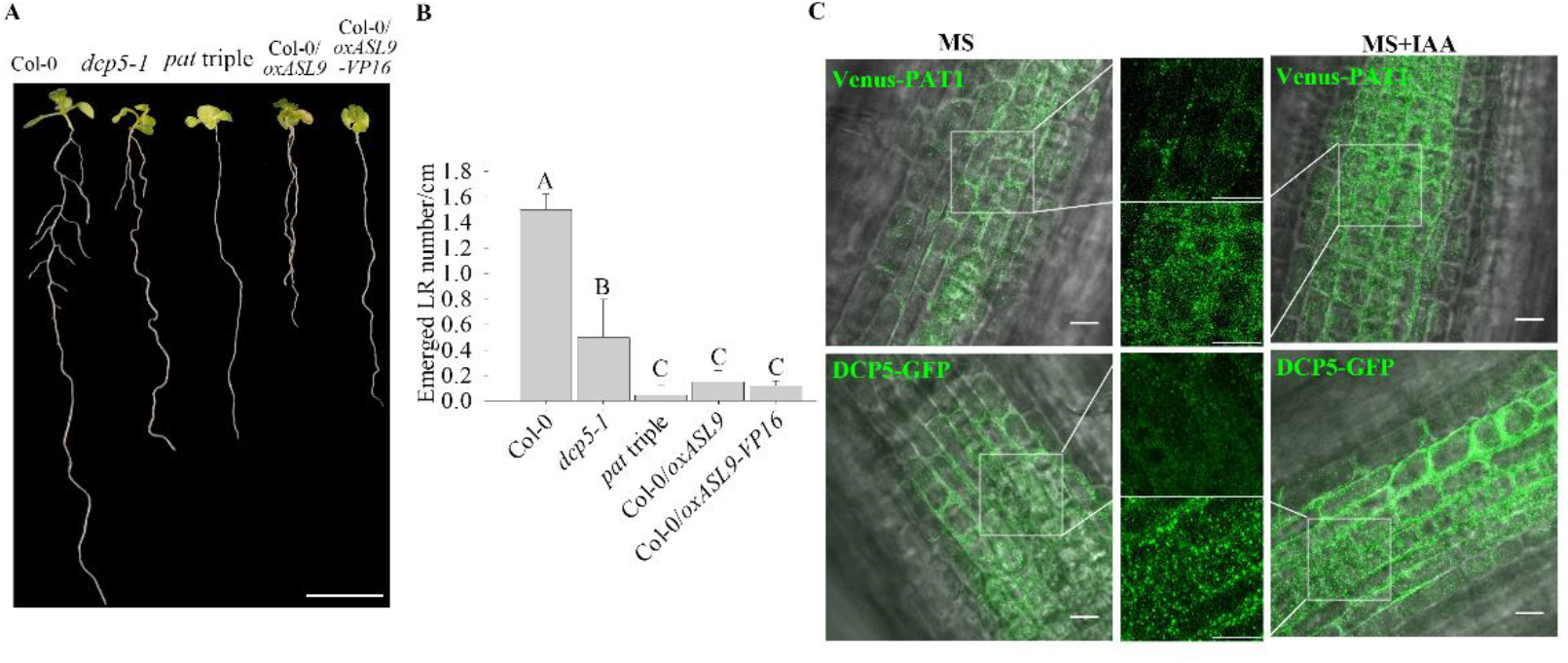
Accumulation of *ASL9* suppresses LR formation. Phenotypes (**A**) and emerged LR density (**B**) of 10-day old seedlings of Col-0, *dcp5-1, pat* triple, Col-0/*oxASL9* and *Col-0/oxASL9-VP16*. The experiment was repeated 4 times, in each repeat sample size (n)>10 for each genotype, and representative pictures are shown. The scale bar indicates 1cm. Bars marked with the same letter are not significantly different from each other (P-value>0.05). (**C**) Representative confocal microscopy pictures of root regions from 7-day old seedlings with either Venus-PAT1 or DCP5-GFP treated with MS or MS+0.2 μIAA for 15min. Scale bars indicate 10 μm.

### *ASL9* directly contributes to apical hooking and LR formation

The overexpression of *ASL9* is sufficient to suppress apical hook and lateral root development. To examine more directly if *ASL9* accumulation contributes to the developmental defects in decapping mutants, we crossed *asl9-1* to both *dcp5-1* and *pat* triple mutant to generate *dcp5-1asl9-1* and *patasl9-1* mutants. However, we only managed to get *dcp5-1asl9-1* due to *ASL9* and *SUMM2* being linked. We then germinated *dcp5-1asl9-1* seedlings in darkness in the presence or absence of ACC and in both conditions *dcp5-1asl9-1* made more stringent hooks than *dcp5-1* but not as tight as Col-0 or *asl9*-1 did, indicating that the loss-of-function of *asl9* can partially suppress *dcp5-1* hookless phenotype (Fig. 4A&B). Moreover, the LR phenotype of *dcp5-1* was also partially restored by mutating *ASL9* (Fig. 4C&D). Thus, our data indicates that *ASL9* contributes to both apical hooking and LR development in mutant with decapping deficiencies.

**Fig 4.**
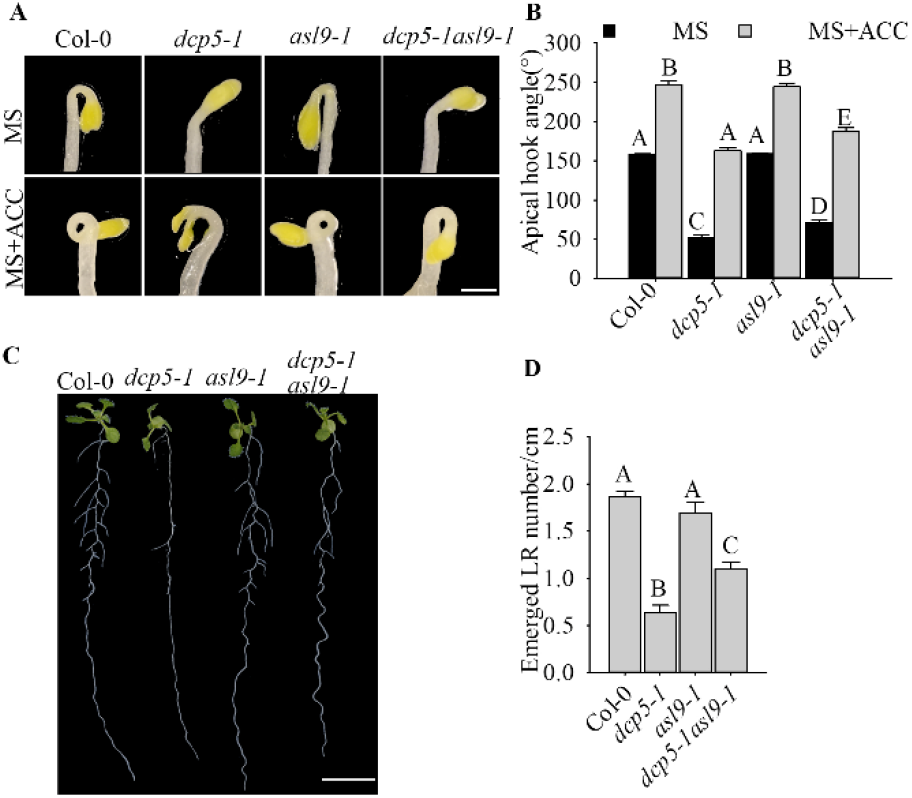
*ASL9* directly contributes to apical hooking and LR formation. Hook phenotypes (**A**) and apical hook angles (**B**) in triple responses to ACC treatment of etiolated Col-0, *dcp5-1, asl9-1* and *dcp5-1asl9-1* seedlings. The treatment was repeated 3 times, in each repeat sample size (n)>50 for each genotype and treatment, and representative pictures are shown. The scale bar indicates 1mm. Phenotypes (**C**) and emerged LR density (**D**) of 10-day old seedlings of Col-0, *dcp5-1, asl9-1* and *dcp5-1asl9-1*. Treatment was repeated 3 times, in each repeat sample size (n)>20 for each genotype, and representative pictures are shown. The scale bar indicates 1cm. Bars marked with the same letter are not significantly different from each other (P-value>0.05).

### Interference with cytokinin signalling and/or exogenous auxin restores developmental defects of *ASL9* over-expressor and mRNA decay deficient mutants

*ASL9* has been implicated in cytokinin signaling (Naito et al., 2007; Ye et al., 2021) in which ARR1, ARR10 and ARR12 are responsible for activation of cytokinin transcriptional responses (Ishida et al., 2008; Xie et al., 2018) and cytokinin acts antagonistically with auxin. Apical hooking and lateral root formation represent classic examples of auxin dependent development (Peer et al., 2011). In support of this, *axr1* mutants showed defective apical hook formation and reduced LR numbers (Estelle and Somerville, 1987;Lehman et al., 1996). We therefore examined cytokinin and auxin related gene expression in both mRNA decay deficient mutants and *ASL9* over-expressor (Fig. S3&4). The cytokinin responsive and signaling repressors type-A ARR genes *ARR8* and *ARR15*, the auxin induced gene *SAUR23* and the auxin biosynthesis gene *TAR2* are all repressed in these genotypes tested, which suggest a misregulation of cytokinin signaling and abrogated auxin homeostasis. To test if the developmental defects of mRNA decay mutants and *Col-0/oxASL9* are due to misregulation of cytokinin, we interfered with cytokinin pathways in *ASL9* over-expressors and decapping mutant *dcp5-1* by knocking out cytokinin signaling activators *ARR10* and *ARR12* (Ishida et al., 2008). Interestingly, both apical hooking and LR formation phenotypes of *ASL9* over-expressors were largely restored in *arr10-5arr12-1* background (Fig. 5) indicating that the developmental defects in *ASL9* over-expressors are most likely caused by misregulation of cytokinin signaling. As for *dcp5-1*, the apical hooking and LR phenotype were partially restored by mutating *arr10* and *arr12* (Fig. 6); which despite not reaching the same extend as seen in *ASL9* over-expressors, was still similar to our observations in *dcp5-1asl9-1* double mutants (Fig 4&5). Furthermore, the expression of *ARR8, ARR15, SAUR23* and *TAR2* in *dcp5-1* were also partially restored in *arr10-5arr12-1* background (Fig. S4). Therefore, our data suggest that apical hooking and LR developmental defects in *ASL9* over-expressors and to some degree in mRNA decapping mutants depend on functional cytokinin signalling.

**Fig 5.**
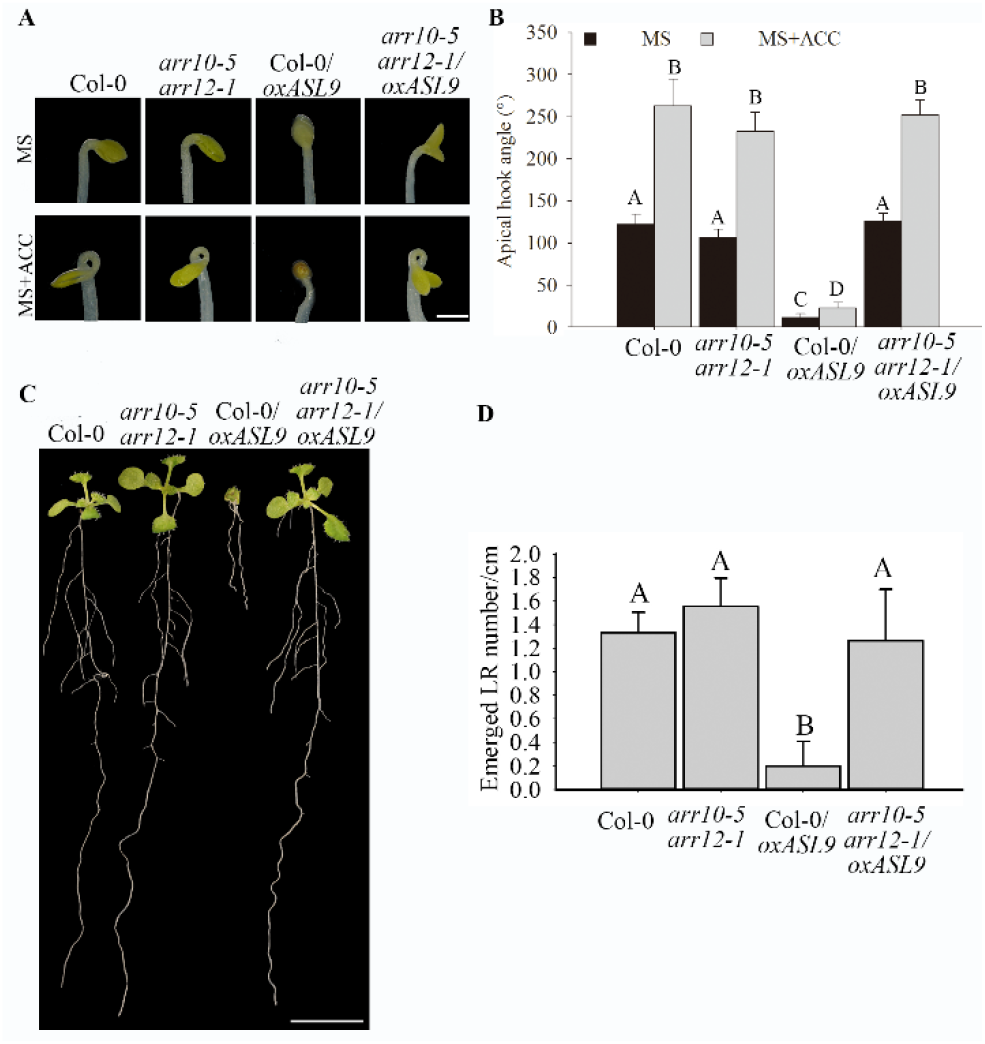
*ARR10* and *ARR12* loss-of-function restores apical hook and LR formation in *ASL9* over-expressor. Hook phenotypes (**A**) and apical hook angles (**B**) in triple responses to ACC treatment of etiolated Col-0, *arr10-5arr12-1*, Col-0*/oxASL9* and *arr10-5arr12-1/oxASL9* seedlings. The treatment was repeated 3 times, in each repeat sample size (n)>20 for each genotype and treatment, and representative pictures are shown. The scale bar indicates 1mm. Phenotypes (**C**) and emerged LR density (**D**) of 10-day old seedlings of Col-0, *arr10-5arr12-1*, Col-0*/oxASL9* and *arr10-5arr12-1/oxASL9*. Treatment was repeated 3 times, in each repeat sample size (n)>10 for each genotype, and representative pictures are shown. The scale bar indicates 1cm. Bars marked with the same letter are not significantly different from each other (P-value>0.05).

**Fig 6.**
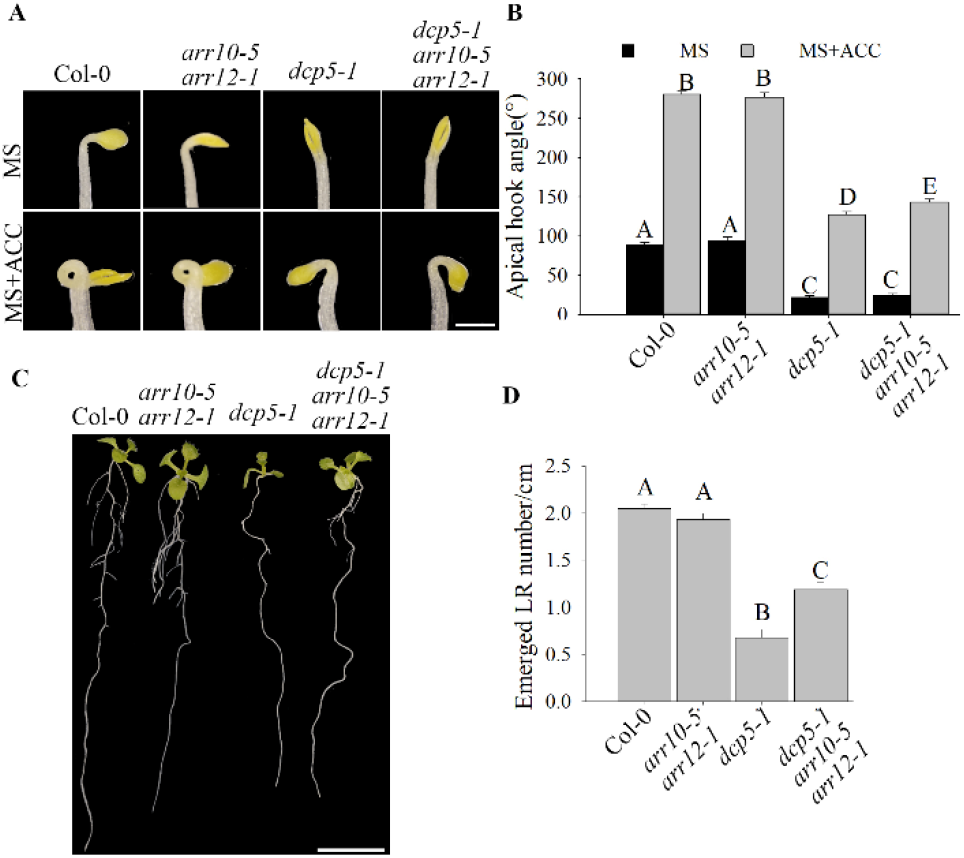
*ARR10* and *ARR12* loss-of-function partially restores apical hook and LR formation in *dcp5-1*. Hook phenotypes (**A**) and apical hook angles (**B**) in triple responses to ACC treatment of etiolated Col-0, *arr10-5arr12-1, dcp5-1* and *dcp5-1arr10-5arr12-1* seedlings. The treatment was repeated 3 times, in each repeat sample size (n)>50 for each genotype and treatment, and representative pictures are shown. The scale bar indicates 1mm. Phenotypes (**C**) and emerged LR density (**D**) of 10-day old seedlings of Col-0, *arr10-5arr12-1, dcp5-1* and *dcp5-1arr10-5arr12-1*. Treatment was repeated 3 times, in each repeat sample size (n)>10 for each genotype, and representative pictures are shown. The scale bar indicates 1cm. Bars marked with the same letter are not significantly different from each other (P-value>0.05).

To test if repressed auxin signaling is also responsible for the developmental defects in mRNA decapping mutants and *ASL9* overexpressors, we firstly confirmed the repressed auxin signaling in mRNA decay mutants by introducing the indirect auxin-responsive reporter *DR5*::GFP. We found increased GFP signals in the concave side of Col-0 apical hook region when dark-grown on MS with/without ACC, but not in *dcp5-1* or *dcp2-1* in either growth condition and the overall GFP signals in *dcp2-1* were markedly lower than Col-0 (Fig. S5). We also examined DR5::GFP signal in the root area of 7-day old Col-0 and *dcp5-1* seedlings and again, overall GFP signal in *dcp5-1* were strikingly lower than Col-0 (Fig. S6). Collectively these data confirmed our supposition that repressed auxin responses in the mRNA decapping mutants affect apical hook and root developmental processes. Consistent with this notion, exogenous auxin supplementation (0.2 μM IAA) lead to partial restoration of LR formation in *dcp5-1, pat* triple and Col-0/*oxASL9* (Fig. S7). Collectively, our findings indicate that misregulation of cytokinin/auxin responses is partially responsible for the developmental defects in the mRNA decay mutants and *ASL9* overexpressors.

## Discussion

Developmental changes require massive overhauls of gene expression (Miyamoto et al., 2015). Apart from unlocking effectors needed to install a new program, previous states or programs also need to be terminated (Tatapudy et al., 2017; Rodriguez et al., 2020). We report here that mRNA decay is required for certain auxin dependent developmental processes. The stunted growth phenotype and downregulation of developmental and auxin responsive mRNAs in the mRNA decapping mutant (Zuo et al., 2022b) supports a model in which defective clearance of mRNAs hampers decision making upon hormonal perception. Apical hooking and LR formation are classic examples of auxin-dependent developmental processes (Peer et al., 2011). In line with this, we and others observed that mRNA decay-deficient mutants are impaired in apical hooking (Fig. 1) and LR formation (Fig. 3) (Perea-Resa et al., 2012; Jang et al., 2019). Interestingly, among the transcripts upregulated in these decay-deficient mutants was that of capped *ASL9/LBD3* (Fig. 2), which is involved in cytokinin signaling (Naito et al., 2007). Cytokinin and auxin can act antagonistically (Su et al., 2011), and cytokinin can both attenuate apical hooking (Tantikanjana et al., 2001) and directly affect LR founder cells to prevent initiation of lateral root primordia (Laplaze et al., 2007). Our findings that defective processing during those developmental events in mRNA decay-deficient mutants involves *ASL9* was supported by our observation that *ASL9* mRNA is directly regulated by the decapping machinery and that Col-0/*oxASL9* transgenic lines cannot reprogram to attain an apical hook or to form LRs (Fig. 2&3) while loss-of-function of *asl9* partially restores the developmental defects in the decapping deficient mutants (Fig. 4). In line with this, we argue that the misregulation of cytokinin-dependent and auxin-dependent signaling is partially responsible for the developmental defects in mRNA decay-deficient mutants. This is supported by the observation that auxin responses in the *dcp5-1* and *dcp2-1* mutants are repressed (Fig. S5&6) and treating *dcp5-1, pat* triple and Col-0/*oxASL9* with exogenous auxin partially restores LR formation (Fig. S7). Besides misregulation of cytokinin signaling pathway in plants overexpressing *ASL9*, short term accumulation of *ASL9* also led to downregulation of cytokinin responsive genes (Ye et al., 2021), indicating a negative role of *ASL9* in regulating cytokinin responses. However, the fact that the developmental defects in *ASL9* over-expressors are largely restored by knocking out 2 cytokinin signaling activator genes *ARR10* and *ARR12* suggest the function of *ASL9* during apical hooking and LR formation largely depends on *ARR10* and *ARR12*. In line with this, the developmental defects of *dcp5-1* are also partially restored in *asl9-1* and *arr10-5arr12-1* backgrounds (Fig. 4&6), but in addition to *ASL9* and *ARR*s, others unidentified factors also contribute to the defects in apical hook and LR formation in decapping mutants.

Arabidopsis contains 42 *LBD/ASL* genes (Matsumura et al., 2009), among these genes *LBD16, LBD17, LBD18* and *LBD29* control lateral roots formation and regulate plant regeneration (Fan et al., 2012) and overexpression of another member *ASL4* also impairs apical hook (Bell et al., 2012). The partial restoration of apical hooking and LR formation caused by *asl9* mutation in *dcp5-1* (Fig. 4) suggest that other *ASL*s and/or non-*ASL* genes also contribute to the developmental defects in decapping mutants. Besides lateral root formation, it was recently reported that *Arabidopsis* LBD3, together with LBD4, functions as rate-limiting components in activating and promoting root secondary growth, which is also tightly regulated by auxin and cytokinin, indicating that LBDs balance primary and secondary root growth (Smetana et al., 2019; Xiao et al., 2020; Smith et al., 2020; Ye et al., 2021). Together with auxin, cytokinin plays crucial roles in vascular development through the two-component signaling system, and plants with mutations in cytokinin receptor or type B *ARRs* exhibit vasculature defects (Mähönen et al., 2006; Kondo et al., 2011; Yokoyama 2007). Hence, we cannot exclude the possibility that the developmental defects we observed in mRNA decapping mutants and *ASL9* over-expressors are also related to their vascular development.

Deadenylated mRNA can be degraded via either 3’-5’ exosomal exonucleases and SUPPRESSOR OF VCS (SOV)/DIS3L2 or via the 5’-3’ exoribonuclease activity of the decapping complex (Garneau et al., 2007; Sorenson et al., 2018). Sorenson et al. (2018) found that *ASL9* expression is dependent on both VCS and SOV based on their transcriptome analysis, so that *ASL9* might be a target of both pathways (Sorenson et al., 2018). While more direct data is needed to conclude whether SOV can directly regulate *ASL9* mRNA levels, we have shown that *ASL9* is a target of the mRNA decapping machinery. However, since the Col-0 accession is a *sov* mutant and has no developmental defects, the SOV decay pathway probably only plays an accessory role in regulation of *ASL9*. The function of PBs in mRNA regulation has been controversial since mRNAs in PBs may be sequestered for degradation or re-enter polysomal translation complexes (Franks and Lykke-Andersen, 2008). Yeast PAT1 has also been found to repress translation (Coller and Parker, 2005) and a recent study has confirmed that PBs function as mRNA reservoirs in dark-grown *Arabidopsis* seedlings (Jang et al., 2019). These data open the possibility that *ASL9* might be also regulated at the translational level by the decapping machinery. Nevertheless, our finding of direct interaction of *ASL9* transcripts with DCP5 and PAT1, together with the accumulation of capped *ASL9* in mRNA decay mutants, indicates that *ASL9* misregulation in *dcp2-1, dcp5-1* and *pat* triple mutants is due to mRNA decapping deficiency (Fig. 2).

## Materials and Methods

### Plant materials and growth conditions

*Arabidopsis thaliana* ecotype Columbia (Col-0) was used as a control. All mutants used in this study are listed in Table S1. T-DNA insertion lines for AT5G13570(*DCP2*) *dcp2-1* (Salk_000519), At1g26110 (*DCP5) dcp5-1* (Salk_008881) and double mutant *arr10-5arr12-1* has been described (Xu et al., 2006; Ishida et al., 2008; Xu and Chua, 2009). The T-DNA line for for AT1g16530 (*ASL9*) is SAIL_659_D08 with insertion in the first exon. Primers for newly described T-DNA lines are provided in Table S2. *pat* triple mutant, Venus-PAT1 and DCP5-GFP transgenic line have also been described (Zuo et al., 2022b; Chicois et al., 2018). The YFP WAVE line was from NASC (Nottingham, UK) (Geldner et al., 2009). Col-0/*oxASL9* line has been described before (Naito et al., 2007).

Plants were grown in 9×9cm or 4×5cm pots at 21°C with 8/16hrs light/dark regime, or on plates containing Murashige–Skoog (MS) salts medium (Duchefa), 1% sucrose and 1% agar with 16/8hr light/dark.

### Plant treatments

For ethylene triple response assays, seeds were plated on normal MS and MS+50μM ACC, vernalized 96hrs and placed in the dark at 21°C for 4 days before pictures were taken. Apical hook angle is defined as 180° minus the angle between the tangential of the apical part with the axis of the lower part of the hypocotyl, in the case of hook exaggeration, 180° plus that angle is defined as the angle of hook curvature (Vandenbussche et al., 2010). Cotyledon and hook regions of etiolated seedlings were collected after placing in the dark at 21°C for 4 days for gene expression and XRN1 assay. For LR formation assays, seeds on MS plates were vernalized 96hrs and grown with 16/8 hrs light/dark at 21°C vertically for 10 days. For external IAA application for LR formation experiments, seeds on MS plates were vernalized 96hrs and grown with 16/8 hrs light/dark at 21°C for 7 days and the seedlings were moved to MS or MS+IAA plates and grown vertically for 7 days.

### Cloning and transgenic lines

pGreenIIM DR5V2-ntdtomato/DR5-n3GFP has been published previously (Liao et al., 2015). Arabidopsis transformation was done by floral dipping (Clough and Bent, 1998) for Col-0/DR5::GFP and thereafter Col-0/DR5::GFP was crossed to *dcp5-1* and *dcp2-1^het^* to achieve *dcp5-1*/DR5::GFP and *dcp2-1/* DR5: :GFP. *arr10-5arr12-1/oxASL9* was generated by vacuum infiltrating *arr10-5arr12-1* with *A. tumefaciens* strain EHA101 harbouring pSK1-ASL9 (Naito et al., 2007). Transformants were selected on hygromycin (30 mg/L) or methotrexate (0.1mg/L) MS agar, and survivors were tested for transcript expression by qRT-PCR and protein expression by immuno-blotting and at least 2 independent lines were used for further analysis.

### Protein extraction, SDS-PAGE and immunoblotting

Tissue was ground in liquid nitrogen and 4×SDS buffer (novex) was added and heated at 95°C for 5 min, cooled to room temperature for 10min, samples were centrifuged 5min at 13000 rpm. Supernatants were separated on 10% SDS-PAGE gels, electroblotted to PVDF membrane (GE Healthcare), blocked in 5% (w/v) milk in TBS-Tween 20 (0.1%, v/v) and incubated 1hr to overnight with primary antibodies (anti-GFP (AMS Biotechnology 1:5.000)). Membranes were washed 3 × 10 min in TBS-T (0.1%) before 1hr incubation in secondary antibodies (anti-rabbit HRP or AP conjugate (Promega; 1: 5.000)). Chemiluminescent substrate (ECL Plus, Pierce) was applied before camera detection. For AP-conjugated primary antibodies, membranes were incubated in NBT/BCIP (Roche) until bands were visible.

### Confocal microscopy

Imaging was done using a Zeiss LSM 700 confocal microscope. The confocal images were analyzed with Zen2012 (Zeiss) and ImageJ software. Representative maximum intensity projection images of 10 Z-stacks per image have been shown in Fig. 1,3, S5 &6.

### RNA extraction and qRT-PCR

Total RNA from tissues was extracted with TRIzol™ Reagent (Thermo Scientific), 2μg total RNA were treated with DNAse I (Thermo Scientific) and reverse transcribed into cDNA using RevertAid First Strand cDNA Synthesis Kit according to the manufacturer’s instructions (Thermo Scientific). The *ACT2* gene was used as an internal control. qPCR analysis was performed on a Bio-RAD CFX96 system with SYBR Green master mix (Thermo Scientific). Primers are listed in Table S2. All experiments were repeated at least three times each in technical triplicates.

### In Vitro XRN1 Susceptibility Assay

Transcripts XRN1 susceptibility was determined as described (Mukherjee et al. 2012; Kiss et al., 2016) with some modification. Total RNA was extracted from tissues using the NucleoSpin^®^ RNA Plant kit (Machery-Nagel). 1μg RNA was incubated with either 1 unit of XRN1 (New England Biolabs) or water for 2hr at 37°C, loss of ribosomal RNA bands on gel electrophoresis was used to ensure XRN1 efficiency, after heating inactivation under 70°C for 10min, half of the digest was then reverse transcribed into random primed cDNA with RevertAid First Strand cDNA Synthesis Kit (Thermo Scientific). Capped target transcript accumulation was measured by comparing transcript levels from XRN1-treated versus mock-treated samples using qPCR (*EIF4A1* serves as inner control) for the individual genotypes (Mukherjee et al. 2012; Roux et al., 2015; Kiss et al., 2016).

### RIP assay

RIP was performed as previously described (Streitner et al., 2012). 1.5g tissues were fixated by vacuum infiltration with 1% formaldehyde for 20min followed by 125 mM glycine for 5min. Tissues were ground in liquid nitrogen and RIP lysis buffer (50mM Tris-HCl pH 7.5; 150mM NaCl; 4mM MgCl2; 0.1% Igepal; 5 mM DTT; 100 U/mL Ribolock (Thermo Scientific); 1 mM PMSF; Protease Inhibitor cocktail (Roche)) was added at 1.5mL/g tissue powder. Following 15 min centrifugation at 4°C and 13000rpm, supernatants were incubated with GFP Trap-A beads (Chromotek) for 4 hours at 4°C. Beads were washed 3 times with buffer (50 mM Tris-HCl pH 7.5; 500 mM NaCl; 4 mM MgCl2; 0.5 % Igepal; 0.5 % Sodium deoxycholate; 0.1 % SDS; 2 M urea; 2 mM DTT before RNA extraction with TRIzol™ Reagent (Thermo Scientific)). Transcript levels in input and IP samples were quantified by qPCR, and levels in IP samples were corrected with their own input values and then normalized to YFP WAVE lines for enrichment.

### 5’-RACE assay

5’-RACE assay was performed using the First Choice RLM-RACE kit (Thermo Scientific) following manufacture’s instruction. RNAs were extracted from 4-day-old etiolated seedlings with the NucleoSpin^®^ RNA Plant kit (Machery-Nagel), and PCRs were performed using a low (26-28) or high (30-32) number of cycles. Specific primers for the 5’ RACE adapter and for the genes tested are listed in Table S2.

### Statistical analysis

Statistical details of experiments are reported in the figures and legends. Systat software was used for data analysis. Statistical significance between groups was determined by one-way ANOVA (analysis of variance) followed by Holm-Sidak test.

---

**Fig S1.**
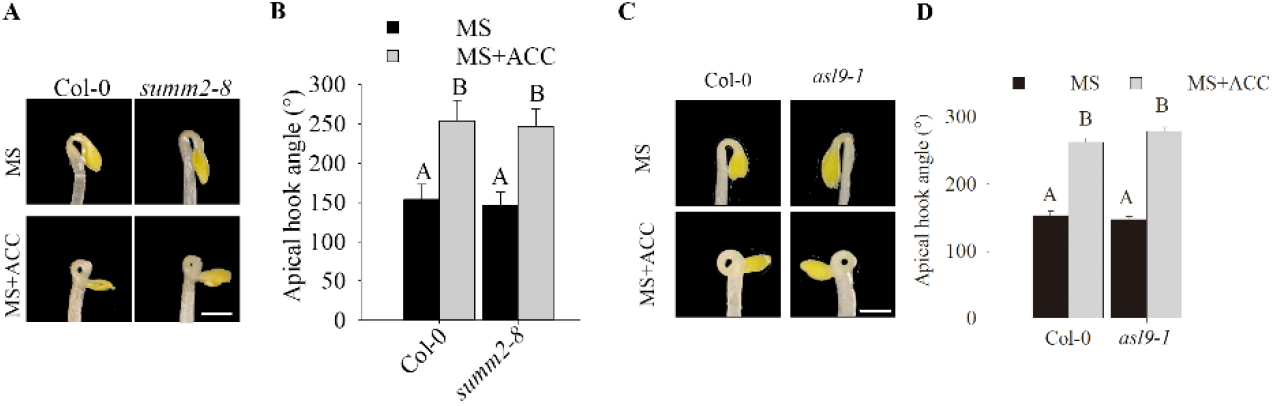
Apical hook in *summ2-8* and *asl9-1* mutants. Hook phenotypes (**A**) and apical hook angles (**B**) in triple responses to ACC treatment of etiolated seedlings of Col-0 and *summ2-8*. Hook phenotypes (**C**) and apical hook angles (**D**) in triple responses to ACC treatment of etiolated Col-0 and *asl9-1* seedlings. The treatment was repeated 3 times(n>30), and representative pictures are shown. The scale bar indicates 1mm. Bars marked with the same letter are not significantly different from each other (P-value>0.05).

**Fig S2.**
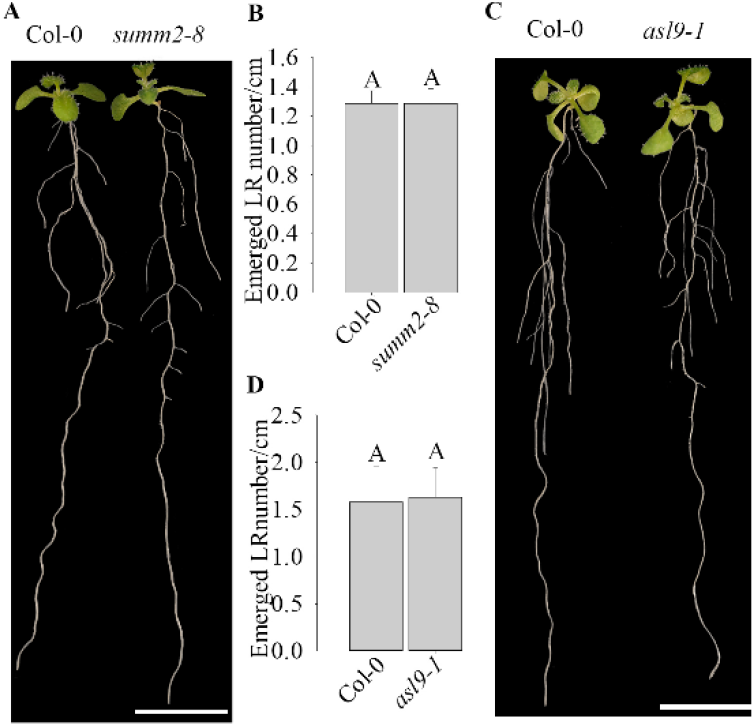
LR formation and primary root growth in *summ2-8* and *asl9-1* mutants. Phenotypes (**A**) emerged LR density and primary root length (**B**)of 10-day old seedlings of Col-0 and *summ2-8*.Phenotypes (**C**) emerged LR density and primary root length (**D**) of 10-day old seedlings of Col-0 and *asl9-1*. The experiment was repeated 3 times(n>10), and representative pictures are shown. The scale bar indicates 1cm. Bars marked with the same letter are not significantly different from each other (P-value>0.05).

**Fig S3.**
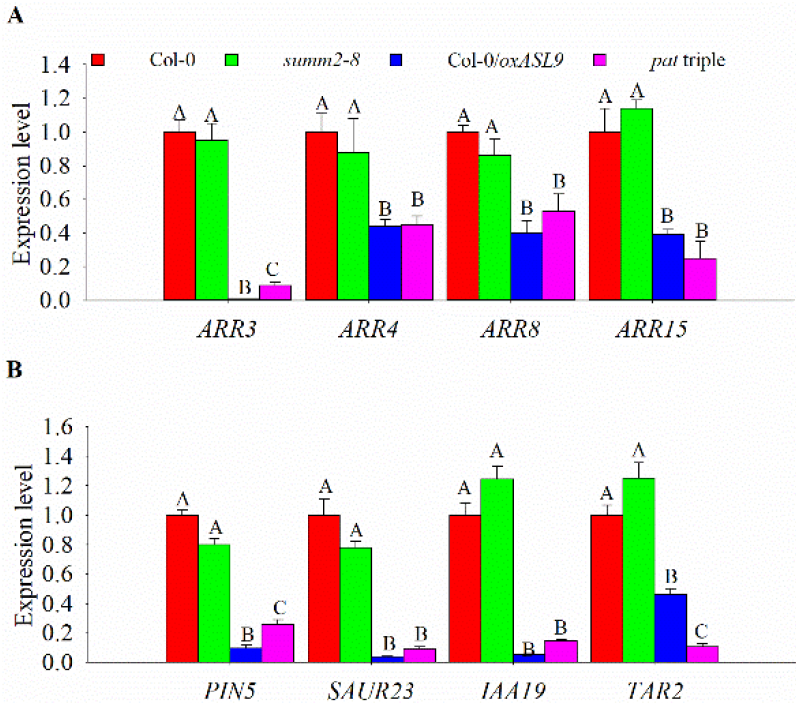
Cytokinin and auxin related genes expression in mRNA decay deficient mutant and *ASL9* over-expressor. Cytokinin pathway repressor genes(**A**) *ARR3, ARR4, ARR8* and *ARR15* and auxin pathway genes(**B**) *PIN5, SAUR23, IAA19 and TAR2* expression levels in 10-day-old seedlings of Col-0, *summ2-8, Col-0/oxASL9* and *pat1-1path1-4path2-1summ2-8*. The experiment was repeated 3 times, error bars indicate SE of bio-triplicates. Bars marked with the same letter are not significantly different from each other (P-value>0.05).

**Fig S4.**
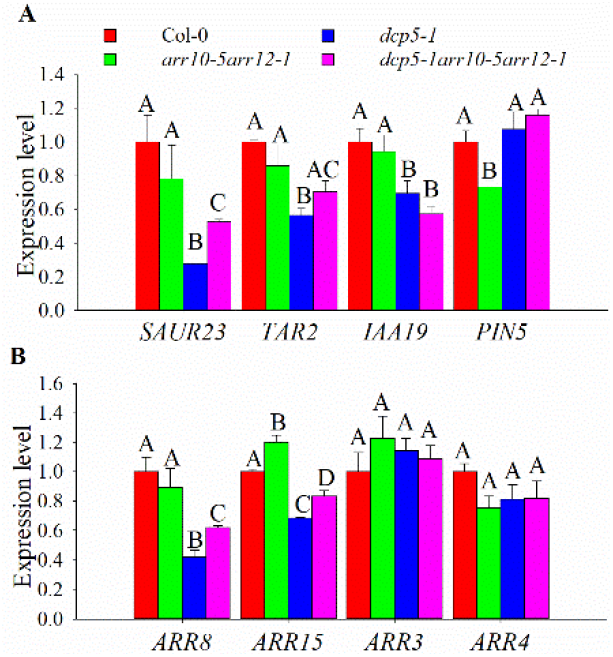
Cytokinin and auxin related genes expression in *dcp5-1* and *dcp5-1arr10-5arr12-1*. Auxin pathway genes(**A**) *SAUR23, TAR2, PIN5 and IAA19* and cytokinin pathway repressor genes(**B**) *ARR8, ARR15, ARR3* and *ARR4* and expression levels in 10-day-old seedlings of Col-0, *arr10-5arr12-1, dcp5-1* and *dcp5-1arr10-5arr12-1*. The experiment was repeated 3 times, error bars indicate SE of bio-triplicates. Bars marked with the same letter are not significantly different from each other (P-value>0.05).

**Fig S5.**
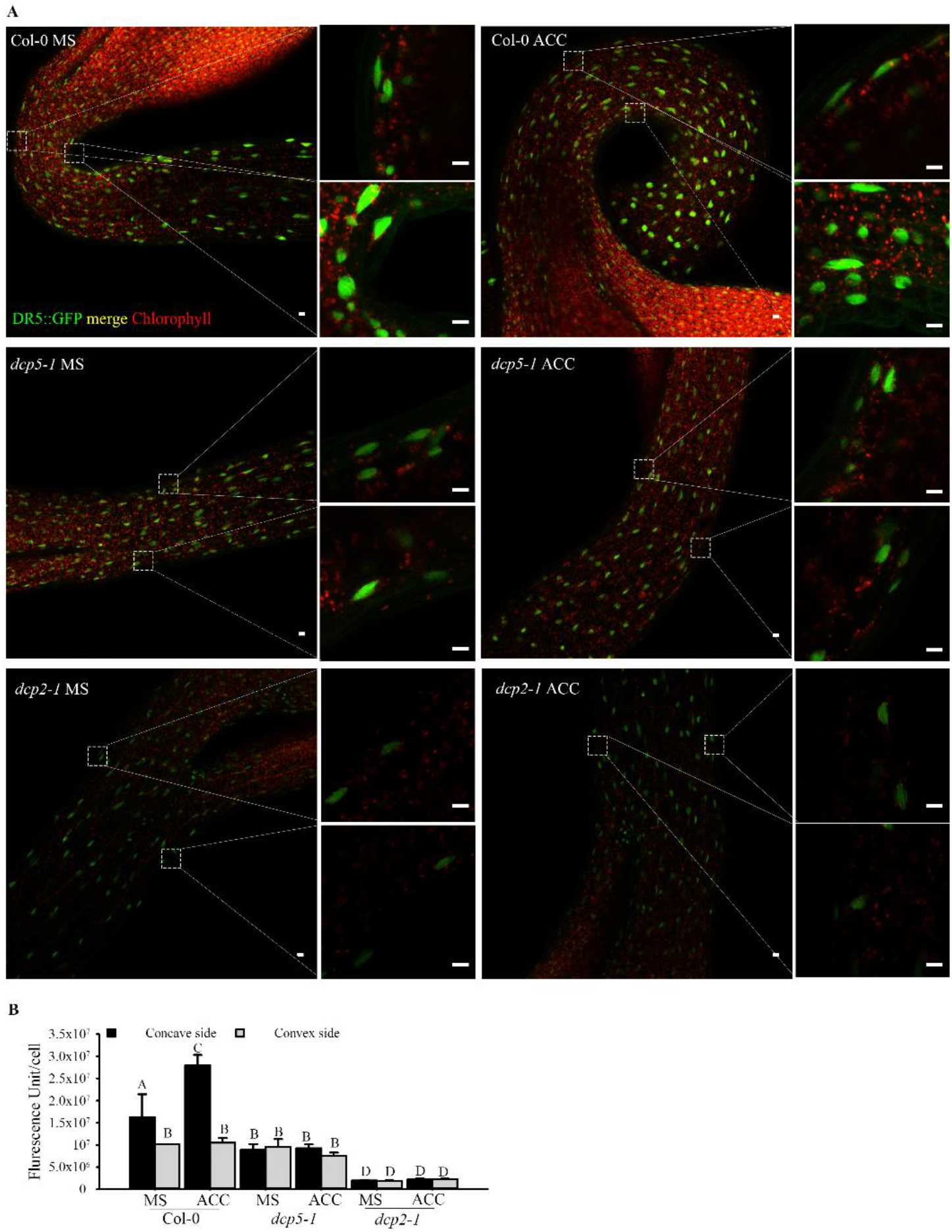
mRNA decapping mutants *dcp5-1* and *dcp2-1* exhibit repressed auxin responses in apical hook regions. Representative confocal microscopy pictures(**A**) and quantification(**B**) of GFP signals in concave and convex side of apical hook regions of Col-0, *dcp5-1* and *dcp2-1* expressed with DR5::GFP following ACC treatment. Seeds of Col-0/DR5::GFP, *dcp5-1*/DR5::GFP and *dcp2-1/* DR5::GFP on MS or MS+ACC plates were vernalized 96hr and grown in dark for 4 days. All pictures were taken under the same confocal microscope settings. Scale bars indicate 10μm. Bars marked with the same letter are not significantly different from each other (P-value>0.05).

**Fig S6.**
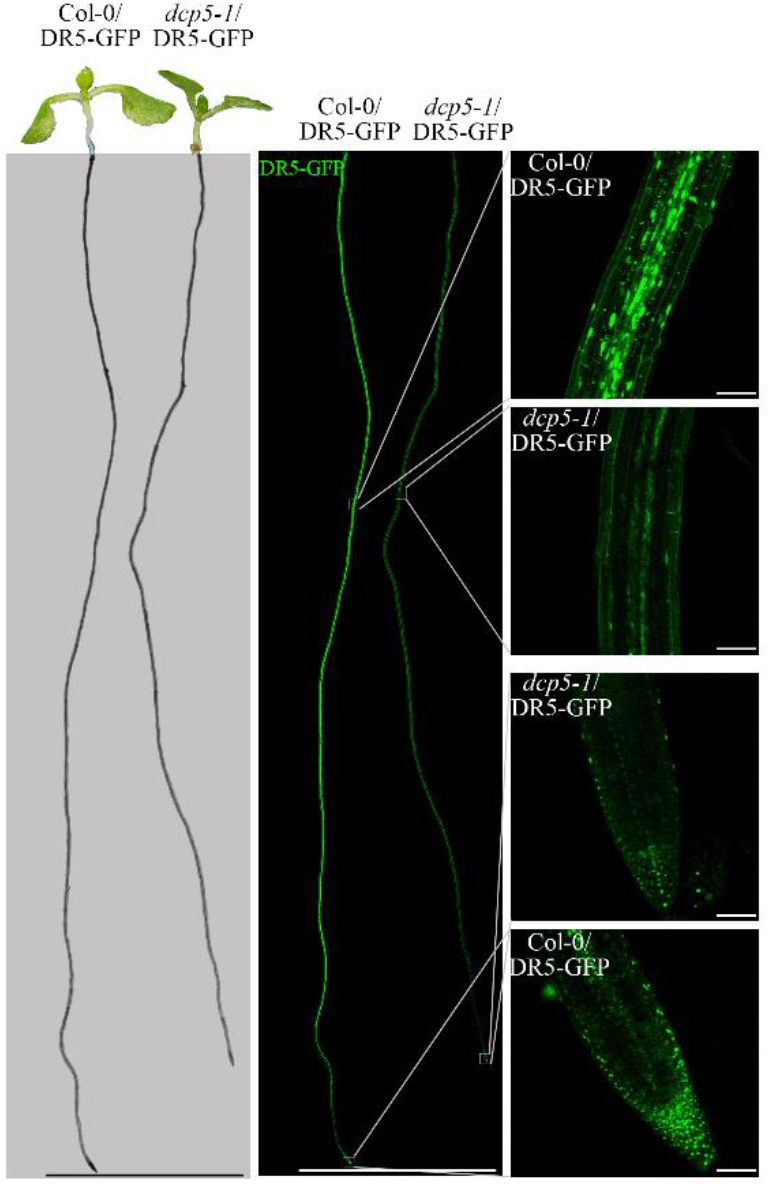
mRNA decapping mutants *dcp5-1* exhibit repressed auxin responses in root regions. Representative confocal microscopy pictures of GFP signals in root regions of 7-day old seedling of Col-0 and *dcp5-1* expressed with DR5::GFP. Seeds of Col-0/DR5::GFP and *dcp5-1*/DR5::GFP on MS plates were vernalized 96hr and grown vertically for 7 days. All pictures were taken under the same confocal microscope settings. Scale bars indicate 1cm in the left and middle lane and100μm.

**Fig S7.**
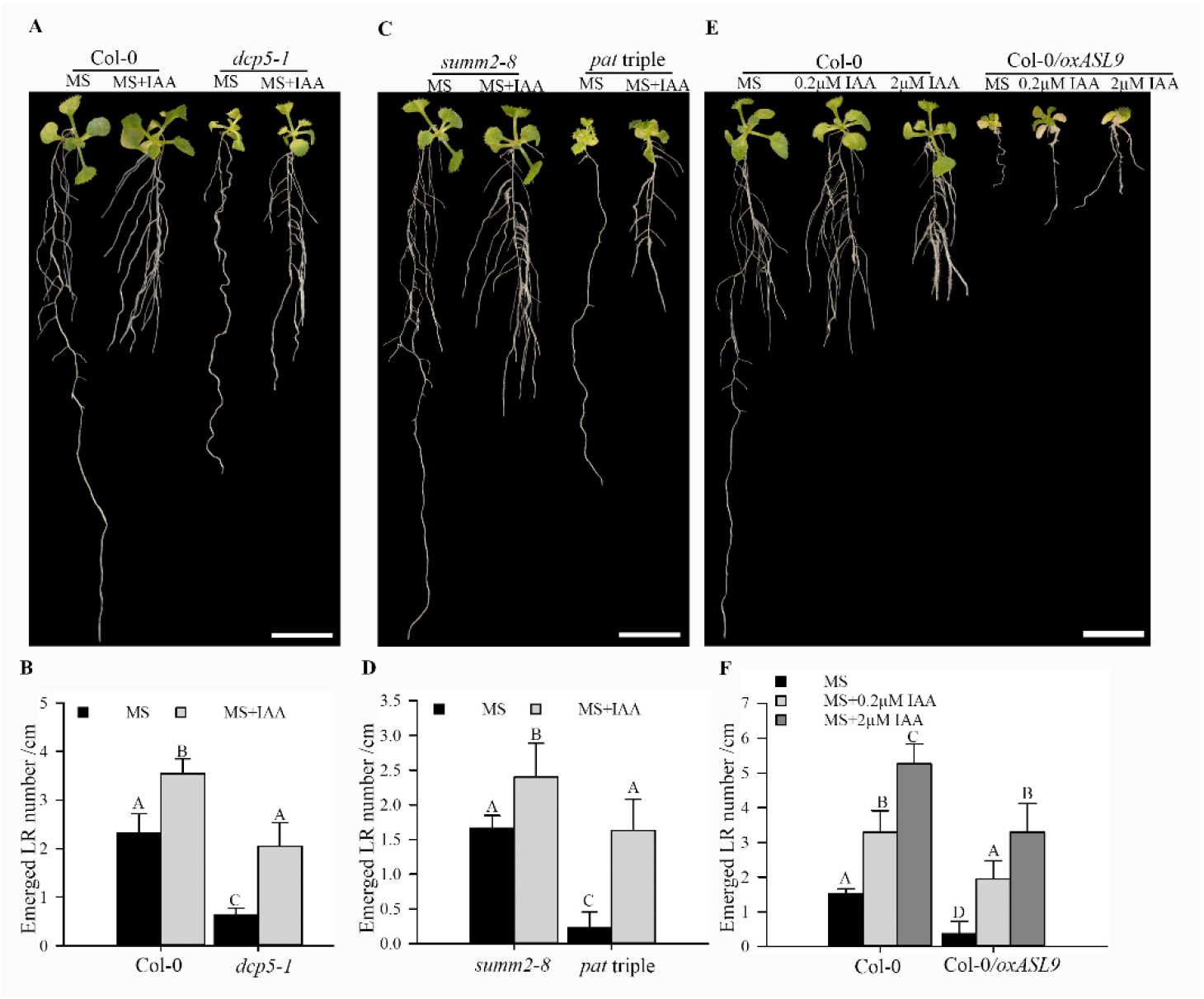
Auxin restores LR formation in mRNA decay deficient mutants and *Col-0/oxASL9*. Phenotypes(**A**) and emerged LR density(**B**) of 14-day old seedlings of *summ2-8* and *pat1-1path1-4path2-1summ2-8* on MS or MS with 0.2μM IAA. Phenotypes (**C**) and emerged LR density (**D**) of 14-day old seedlings of Col-0 and *dcp5-1* on MS or MS with 0.2μM IAA. Phenotypes (**E**) and emerged LR density (**F**) of 14-day old seedlings of Col-0 and Col-0/*oxASL9* on MS, MS with 0.2μM IAA or MS with 2μM IAA. Seeds on MS plates were vernalized 96hrs and grown with 16/8 hrs light/dark at 21°C for 7 days. The seedlings were moved to MS or MS+IAA plates and grown vertically for 7 days. The treatment was repeated 3 times, sample size (n)>10 for each genotype and treatment, and representative pictures are shown. The scale bar indicates 1cm. Bars marked with the same letter are not significantly different from each other (P-value>0.05).

## Acknowledgments

We thank Qi-Jun Chen for Phee401, Nam-Hai Chua for *dcp5-1* and *dcp2-1* seeds and Damien Garcia for DCP5-GFP marker line seeds. Special thanks to John Mundy for advice throughout the project and critically reading the manuscript. This work was supported by the Novo Nordisk Fonden and the Hartmanns Fond to MP (NNF18OC0052967 and A32638), the Danish Research Agency grant to ER (DFF1-1032-00249B), the Institute Strategic Programme grant (BB/P013511/1) to the John Innes Centre and a PhD scholarship from China Scholarship Council to ZZ (201504910714). Microscopy was performed at the Center for Advanced Bioimaging, University of Copenhagen.

ZZ, MER, and MP conceived and designed the experiments. ZZ, MER, JRC, YD, TY, SDH, EK, LØ and ER performed experiments. ZZ and MP analyzed the data. ZZ, ER and MP wrote the manuscript.

The authors declare no competing interests.

Correspondence and requests for materials should be addressed to MP.

