## Supplementary material for "The mRNA decapping machinery targets *LBD3/ASL9* to mediate apical hook and lateral root development in *Arabidopsis*": Table S1

**Table S1. Mutants used in this study**

| Mutant | Information | Source |
|---|---|---|
| *path1-4* | 7bp deletion in Exon 2, frame shift and early stop codon | Zuo et al., 2022b |
| *path2-1* | 35bp deletion in Exon 2, frame shift, early stop codon | Zuo et al., 2022b |
| *asl9-1* | SAIL_659_D08, T-DNA insertion in Exon 1 | NASC (Nottingham, UK) |
| *pat1-1* | Salk_040660, T-DNA insertion in Exon 5 | Roux et al., 2015 |
| *summ2-8* | SAIL_1152A06, T-DNA insertion in Exon 1 | Zhang et al., 2012 |
| *dcp5-1* | Salk_008881, T-DNA insertion in 3'-UTR | Xu and Chua, 2009 |
| *dcp2-1* | Salk_000519, T-DNA insertion in Exon 3 | Xu et al., 2006 |
| *arr10-5* | Salk_098604, T-DNA insertion in Exon 5 | Ishida et al., 2008 |
| *arr12-1* | Salk_054752, T-DNA insertion in Exon 3 | Ishida et al., 2008 |

**Table S2. Primers used in this study**

| Primer | Sequence | Use |
|---|---|---|
| **genotyping** | | |
| LBb1.3 | ATTTTGCCGATTTCGGAAC | Genotyping Salk lines |
| SAIL LB3 | CATCTGAATTTCATAACCAATCTC | Genotyping Sail lines |
| SAIL_659_D08 LP | ATGTTGTACGTTGATTTGGGG | SAIL_659_D08 genotyping |
| SAIL_659_D08 RP | TATTCTTTACACGCGGTTTCG | SAIL_659_D08 genotyping |
| DCP2LP | TGATGGGGTTTTGTTTCAGTC | *dcp2* genotyping |
| DCP2RP | ACTATGATCAATGAGTGGCGG | *dcp2* genotyping |
| **qPCR** | | |
| ASL9 for | CAAAAGGGTCACAGACACGGAA | qPCR of *ASL9* |
| ASL9 rev | GGCCTCGTACACCATCGAATC | qPCR of *ASL9* |
| EIF4A1F | GATCTGCACCAGAAGGCACA | qPCR of *EIF4A1* |
| EIF4A1R | CCCAGTACCAGACTGAGCCTGTTG | qPCR of *EIF4A1* |
| ARR3F | GAAACTCGCCGACGTGAAAC | qPCR of *ARR3* |
| ARR3R | TCCACAAGCGAAGTTGCAGA | qPCR of *ARR3* |
| ARR4F | ATGGCCAGAGACGGTGGTGTTTC | qPCR of *ARR4* |
| ARR4R | ATCTAATCCGGGACTCCTCATC | qPCR of *ARR4* |
| ARR8F | GACCCAAATGCACTCTCTACATC | qPCR of *ARR8* |
| ARR8R | CTCTTCAGCTCCTTCTTCCAAAC | qPCR of *ARR8* |
| ARR15F | GACGACTGTTGAGAGTGGGAC | qPCR of *ARR15* |
| ARR15R | CTCCTCTGCTCCTTCTATCATAC | qPCR of *ARR15* |
| PIN5-FW | CCATCGGCTCTATTGTCCTTG | qPCR of *PIN5* |
| PIN5-RV | GCGACGAGCACAGGTAGAGA | qPCR of *PIN5* |
| SAUR23 F | ATTCAAACTTTCAGACAAAAGAAATGG | qPCR of *SAUR23* |
| SAUR23 R | ACAAGGAAACAACTCTATCTCTAACT | qPCR of *SAUR23* |
| IAA19 F | GGTGACAACTGCGAATACGTTACCA | qPCR of *IAA19* |
| IAA19 R | CCCGGTAGCATCCGATCTTTTCA | qPCR of *IAA19* |
| TAR2 F | CATGATTTGGCTTACTATTGGCCACAG | qPCR of *TAR2* |
| TAR2 R | GTCTTTCACCAAAGCCCATCCAATC | qPCR of *TAR2* |
| ARR10F | GCTTCTGATGCTGGTTCCTT | qPCR of *ARR10* |
| ARR10R | CAATCACCTTCCGAGAAATCA | qPCR of *ARR10* |
| ARR12F | CTCCACGATGAAGCAGGAA | qPCR of *ARR12* |
| ARR12R | AACTAAACCCTCCATATCCCAAA | qPCR of *ARR12* |
| **5'-RACE** | | |
| ASL9 inner | GGCCTCGTACACCATCGAATC | RACE inner PCR |
| ASL9 outer | ATGTCGATGTCACTGTAGAAG | RACE outer PCR |
| EIF4A1 inner | GGTTCTCTTGAAGACCCATGGCATC | RACE inner PCR |
| EIF4A1 outer | CCCAGTACCAGACTGAGCCTGTTG | RACE outer PCR |
